# Comparative tissue-specific transcriptomics reveal the genetic bases underlying evolutionary convergence of seed shattering in two independently evolved weedy rice lineages

**DOI:** 10.1101/2024.09.28.615632

**Authors:** Xiang Li, Ana L. Caicedo

## Abstract

The repeated evolution of high seed shattering during multiple independent de-domestications of cultivated Asian rice (*Oryza sativa*) into weedy rice is a prime example of convergent evolution. Weedy rice populations converge in histological features of the abscission zone (AZ), a crucial structure for seed abscission, while ancestral cultivated rice populations exhibit varied AZ morphology and levels of shattering. However, the genetic bases of these phenotypic patterns remain unclear. We examined the expression profiles of the AZ region and its surrounding tissues at three developmental stages in two low-shattering cultivars of *aus* and *temperate japonica* domesticated varieties and in two genotypes of their derived high-shattering weed groups, Blackhull Awned (BHA) and Spanish Weedy Rice (SWR), respectively. Consistent with the greater alteration of AZ morphology during the de-domestication of SWR than BHA, fewer genes exhibited a comparable AZ-region exclusive expression pattern between weed and crop in the *japonica* lineage than in *aus* lineage. Transcription factors related to the repression of lignin and secondary cell deposition, such as, *OsWRKY102* and *OsXND-1-like*, along with certain known shattering genes involved in AZ formation, likely played a role in maintaining AZ region identity in both lineages. Meanwhile, most genes exhibiting AZ-region exclusive expression patterns do not overlap between the two lineages and the genes exhibiting differential expression in the AZ region between weed and crop across the two lineages are enriched for different gene ontology terms. Our findings suggest genetic flexibility in shaping AZ morphology, while genetic constraints on AZ identity determination in these two lineages.

**Significance statement:** Exploring the extent of genetic convergence that underlies the morphological convergence - specifically, the recurrent evolution of complete abscission zones in independently evolved weedy rice populations originating from different cultivated rice populations with varying degrees of disrupted abscission zones - can improve our understanding not only of the genetic mechanisms behind convergent evolution, but also of the genetics underlying the agriculturally importance trait of seed shattering.

## Introduction

Evolutionary convergence refers to phenotypic similarities that have arisen in independently evolved species or populations. Convergent evolution is thought to be an adaptive process, where organisms respond similarly to the same environmental challenges. Because it comprises naturally occurring evolutionary replicates, convergent evolution serves as a valuable framework for understanding the flexibility or constraints governing the process of adaptation (Kitano *et al*., 2022; Futuyma, 2010; Losos, 2011; Orr, 2005; Sackton and Clark, 2019). An open question in biology is whether observed phenotypic convergence is often associated with convergence at the genetic level, and, if so, under what circumstances and to what extent these associations exist (Sackton and Clark, 2019).

In the context of agriculture, abscission, the process by which plants shed organs at a given developmental stage, is a trait that has been consistently altered in response to various environmental settings. The purpose of abscission is to get rid of unnecessary organs, like senescent leaves, or to disperse seeds or fruits. Natural seed shattering in wild plants is an abscission process of great ecological importance for the dispersal of mature seeds and the propagation of future generations (Dong and Wang, 2015). However, this trait is not favored in many cultivated plants, particularly cereals, and has been selected against to facilitate harvesting and to avoid yield loss (Fuller, 2007; Fuller and Allaby, 2009). In contrast, weeds in agricultural fields often leverage the shattering trait to improve their survival and maximize their dispersal (Vigueira *et al*., 2013; Maity *et al*., 2021). Thus, abscission has constantly been a target of selection in different groups within the agricultural setting: a reduced shattering during domestication and an elevated shattering level during weed evolution.

Rice (*Oryza sativa*) is an excellent system to examine the convergence of abscission, as seed shattering has been repeatedly modified throughout its evolutionary history. Stemming from two separate but entangled domestication events (Jing *et al*., 2023; Civáň *et al*., 2015; Choi *et al*., 2017), five major rice cultivar subgroups, *aus* (also referred to as circum-Aus), *indica*, *temperate japonica*, *tropical japonica*, and *aromatic* (also referred to as circum-Basmati) are recognized today (Garris *et al*., 2005; Wang *et al*., 2018), and they are all characterized by reduced seed shattering levels compared with the wild ancestor of rice (*Oryza rufipogon*/*nivara*) (Thurber *et al*., 2010). Weedy rice (*Oryza* spp.) is a related weed of cultivated rice and aggressively outcompetes cultivated rice in crop fields. Weedy rice has evolved multiple times independently, most often from crop ancestors through a process of de-domestication (Cao *et al*., 2006; Li *et al*., 2017; Li *et al*., 2022; Imaizumi *et al*., 2021; Hoyos *et al*., 2020; Qiu *et al*., 2020; Reagon *et al*., 2010), each time gaining a competitive advantage through re-acquisition of high seed shattering (Thurber *et al*., 2010; Li *et al*., 2022; Huang *et al*., 2017). The convergence of reduced shattering among different rice cultivars and of regained shattering in various weedy rice populations can provide a robust framework for examining convergent evolution of the abscission process.

We have previously investigated the convergence of seed shattering in rice at the morphological level through examination of the abscission zone (AZ), a specialized cell layer responsible for seed detachment (Li *et al*., 2024). In rice, the AZ is positioned in the rachilla, above rudimental glumes and below sterile lemma. The AZ-type cells are typically small, cytoplasmically dense, and lack lignin compared with their surrounding cells (Li *et al*., 2024). We have observed morphological convergence among independently evolved weedy rice groups, with all weedy rice displaying a complete and obvious AZ (Li *et al*., 2024). In contrast, distinct AZ morphologies with different levels of disruption are prevalent in different cultivated rice populations (Li *et al*., 2024). In particular, the number and distribution of AZ-type cells varies greatly and significantly impacts seed shattering level (Li *et al*., 2024). However, the extent to which there is convergence in the genetic mechanisms governing AZ-type cell identity, cell number, cell distribution, and, ultimately, shattering levels in different rice lineages is unknown.

Here, we conducted a comparative tissue-specific RNA-seq analysis at three developmental stages for two cultivated and two weedy rice genotypes. We selected two cultivated rice varieties stemming from different domestication events that differ drastically in AZ morphology, *aus* and *temperate japonica*, and their de-domesticated weedy rice descendants, BlackHull Awned (BHA) and Spanish Weedy Rice (SWR) respectively (Figure 1), that converge in both high seed shattering and in AZ morphology. Each crop-weed pair thus represents two independent evolutionary lineages, the *aus* lineage and the *japonica* lineage, in which seed shattering and the AZ have been modified. We also examined AZ morphologies across our sampled developmental stages and their correspondence to patterns of gene expression. The AZ-type cell characteristics of lack of lignin, small size, and thin cell walls were consistently observed throughout development. Transcription factors related to the repression of lignin and secondary cell deposition, along with certain known shattering genes involved in AZ formation, likely played a role in maintaining AZ region identity in both lineages. However, the majority of genes correlating with the morphological differences in the AZ between weedy and cultivated rice were unique to each lineage. More genes exhibited shared AZ-region exclusive expression pattern in domesticated and weedy genotypes in the *aus* lineage than in the *japonica* lineage; and the modules exhibiting an AZ-region exclusive expression pattern in the weed but no expression in the crop were only found in the *japonica* lineage. These findings aligned with the higher level of AZ morphological similarity between BHA and *aus* compared to SWR and *temperate japonica*. Our study reveals the degree of genetic convergence and divergence shaping AZ morphology, enhancing our understanding of the evolutionary malleability of the abscission process under selection.

**Figure 1.**
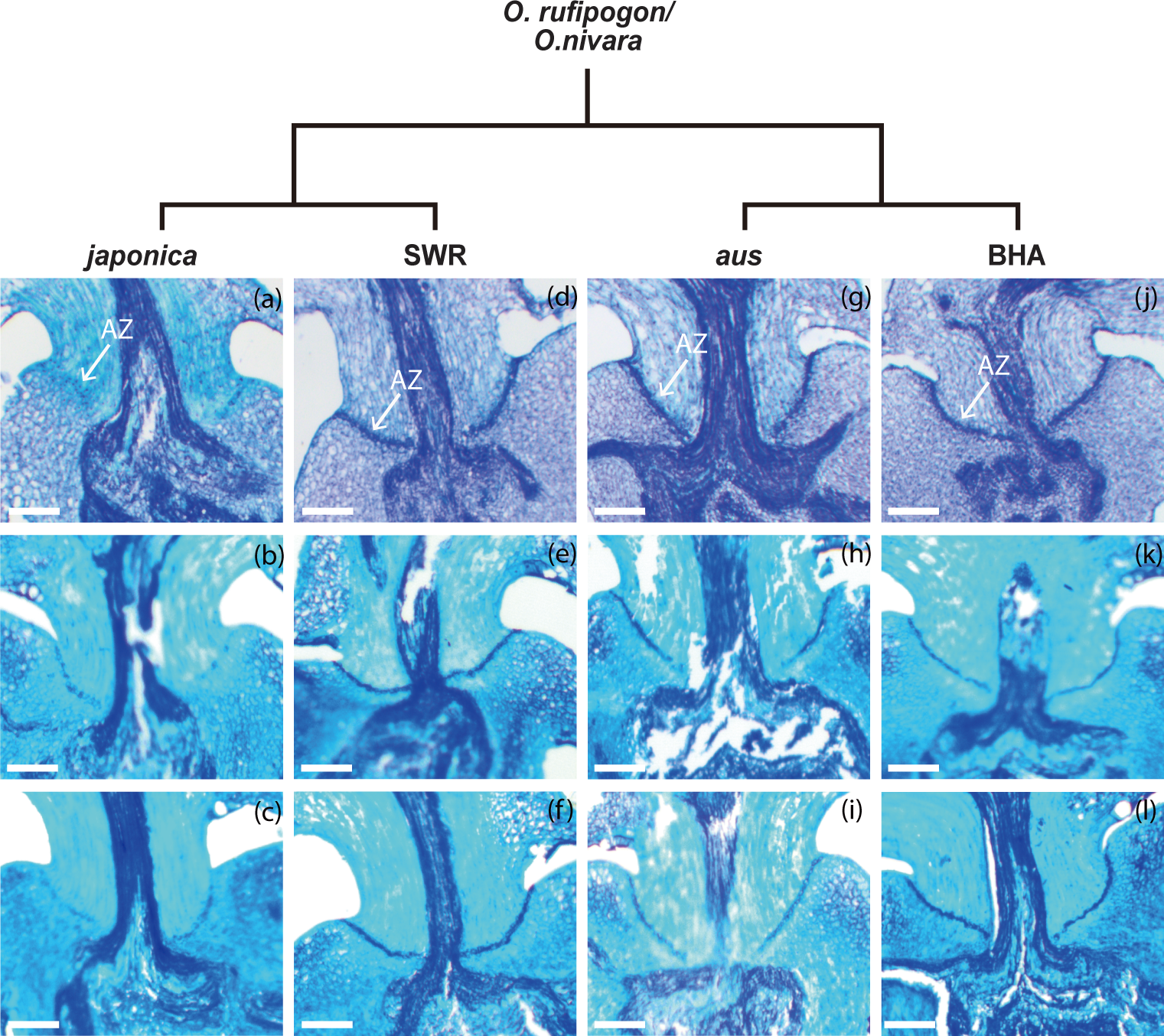
AZ morphology in two lineages of cultivated and weedy rice across development. The phylogeny depicts the known relationship between each weedy and cultivated rice group included in the study. (a-l) Toluidine blue stained rice florets; the AZ cell is stained dark blue, and the AZ region is indicated by the white arrow. Bars: 100 *μm*. (a-c). AN-09-03, a *japonica* crop from BFT, FT to LFT; (d-f). ASF-09-54, a SWR weed from BFT, FT to LFT; (g-i). sau7, an *aus* crop from BFT, FT to LFT; and (i-l) rr14, a BHA weed from BFT, FT to LFT.

## Results

### Alterations in seed shattering levels and AZ morphology across development in weedy and cultivated rice

We have previously established that seed shattering levels in rice typically stabilize at 30 days after heading, when the panicle has emerged halfway from the boot (Thurber *et al*., 2010; Thurber *et al*., 2011). Using Breaking tensile strength (BTS), a widely employed measure for assessing the force required to separate florets or seeds from plants (Turra *et al*., 2023), we examined if each of our weedy and cultivated genotypes of the *aus* and *japonica* lineages followed this typical shattering trajectory throughout development. We measured BTS at heading/flowering time (FT), one week after heading (LFT), and 30 days after heading (LFT30). We attempted to measure BTS one week prior to heading (BFT) but failed due to the fragility of floral tissue. Large variation was evident in all four genotypes during the two early developmental stages (Table S1) with no significant increase or decrease of BTSs from FT to LFT in the *japonica* lineage while relatively clear decrease from FT to LFT in the *aus* lineage (Figure S1). At LFT30, BTS decreased for all genotypes with values consistent with their expected seed shattering abilities. At this point, the weedy BHA and SWR genotypes exhibited high shattering tendencies with median BTS values of 0 gf (Table S1, Figure S1); the *aus* genotype, which is known to belong to a relatively easy-threshing cultivated group, presented a median BTS of 5.50 gf; and the *temperate japonica* genotype, which belongs to a generally low shattering cultivated variety, had an LFT30 median BTS of 87.8 gf, distinctive from the other genotypes (Table S1, Figure S1).

To understand the relationship between shattering levels and AZ morphology throughout development, we also examined the histology of AZs in these four genotypes at three developmental stages: BFT, FT and LFT. Longitudinal sections on seeds collected at LFT30 cannot be conducted due to increased starch content and hardness of the seeds, and the seeds being prone to separation from the pedicel in high shattering rice. In the two high shattering weedy rice genotypes and in the relatively easy shattering *aus* cultivar, the AZ cell layer was visibly present throughout all three developmental stages (Figure 1d-1l). The AZ-type cells distinguished themselves from neighboring cells with their distinctive dark purple color, indicating a lack of lignification, thin walls, and smaller sizes. A small gap existed between the AZ cell layer and middle vascular bundle in the *aus* cultivar (Figure 1g-1i), serving as the major difference in AZ morphology between BHA and *aus* and corresponding to their shattering level disparities. In contrast, in the AZ region of the *temperate japonica* cultivar, no distinct AZ cell layer formed at any developmental stage and only a few scattered AZ-type cells were present (Figure 1a-1c). Non-AZ-type cells within the AZ region exhibited a turquoise color, suggesting lignification. Additionally, these non-AZ-type cells featured thick rather than thin cell walls compared to AZ-type cells, underscoring their cellular distinctions.

### Effects of developmental stage, lineage, and tissue type on gene expression patterns

To identify the genes contributing to AZ development in these four rice genotypes, we conducted tissue-specific RNA-seq analyses. This involved the manual dissection of the AZ region, of the adjacent tissue toward floret side, and of the adjacent tissue toward pedicel side, at the developmental stages corresponding to our examination of AZ histology: BFT, FT and LFT. For each genotype, we collected three biological replicates of each tissue at each developmental stage. The RNA-seq reads were mapped to an *indica* reference genome, R498 (Du *et al*., 2017) with an average overall mapping rate greater than 85%. Principal component analyses (PCA) indicated that developmental stage played the most significant role in distinguishing the expression profiles among samples (Figure 2a, Table S2). Notably, the early developmental stage was clearly separated from the two later developmental stages along principal component 1 (PC1), explaining 62.3% of the variation. The genetic background specific to the evolutionary lineage was the second factor shaping the clustering of the expression patterns (Figure 2b). Along PC2, which accounts for 30.7% of the variation, a clear separation between the *aus* and *japonica* lineages was evident. However, the distinction between weedy and cultivated varieties within the same lineage was not as pronounced as the separation by lineages, despite their shattering differences. Tissue specific expression patterns were observed, with replicates from the same tissues clustering together, particularly in the latter developmental stages (Figure 2c). These patterns also exhibited variation between different lineages. During the early developmental stage, the separation among tissues was more apparent in the *aus* lineage compared to the *japonica* lineage (Figure 2c).

**Figure 2.**
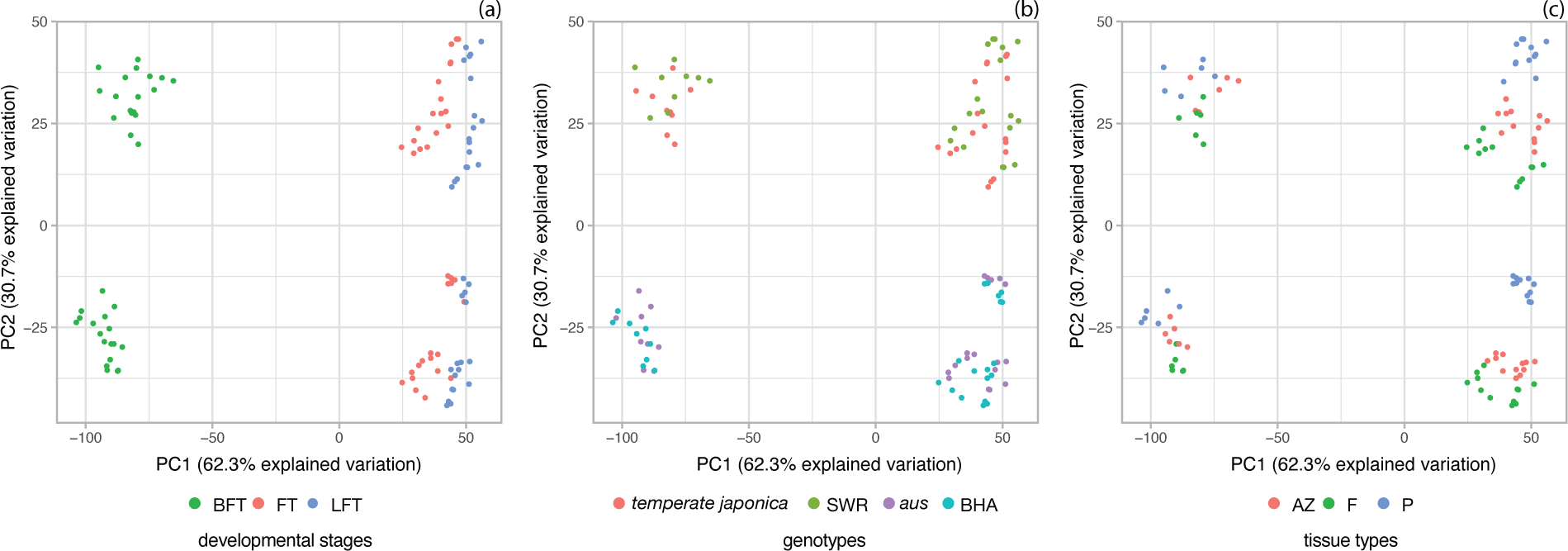
Patterns of gene expression across tissues and developmental stages. (a-c). Principal Component Analysis (PCA) of gene expression profiles of samples collected from three tissue types, abscission zone (AZ), tissue toward floret size (F) and tissue toward pedicel side (P), in four genotypes, *aus*, *temperate japonica*, BHA and SWR, at three developmental stages, BFT, FT and LFT. (a) samples are labelled by different developmental stages; (b) samples are labelled by four genotypes; and (c) samples are labelled by distinct tissue types.

### AZ-region associated modules and their relation to AZ morphology in weedy and cultivated rice from each lineage

Modules represent groups of genes that share similar expression profiles, often indicative of functional relatedness and co-regulation (Saelens *et al*., 2018). We employed Weighted Gene Co-expression Network Analysis (WGCNA) (Langfelder and Horvath, 2008) to identify modules exhibiting exclusive expression in the AZ region, reasoning that genes within these modules are more likely to affect seed shattering abilities. We performed module detection by jointly analyzing the expression profiles of the cultivar and its corresponding weedy counterpart. In the WGCNA analyses for the *temperate japonica* and SWR, we used gene counts obtained from alignment to a *japonica* reference genome (Kawahara *et al*., 2013) to mitigate potential issues arising from genetic disparities between *japonica* and *indica*, which could lead to missing sequence or misalignment (Li *et al*., 2023).

We first clustered samples from each lineage using their expression profiles. The results mirrored the patterns observed in PCA, clearly distinguishing samples collected in the early developmental stage from those in the two latter stages (Figure S2 and S3). During both the BFT and LFT stages, AZ samples from the *aus* and BHA genotypes consistently grouped into a single sub-cluster (Figure S2), suggesting the presence of a similar expression pattern, aligning with their similar AZ histology (Figure 1g-1l). In contrast, the AZ samples in the *japonica* lineage usually clustered with the pedicel or floret samples of the same genotype at each given developmental stage, indicated distinct expression profiles in the AZ regions of *temperate japonica* and SWR (Figure S3). The notable differences in the AZ histology between these two cultivated and weedy g*e*notypes could account for this disparity (Figure 1a-1f).

### Modules explaining AZ similarities and differences between weedy and cultivated rice

We identified modules displaying an AZ-region exclusive expression pattern in both weedy and cultivated genotypes from the same lineage. These modules are more likely to contribute to the AZ region identity or to AZ morphological features shared between genotypes within a lineage. We correlated the eigengene expression, a representative summary measure of the gene expression profiles within a module, with tissue classification by assigning 1 to samples from the AZ region in both cultivated and weedy genotypes and 0 to all other tissues. Modules with a correlation co-efficiency greater than 0.7 and a p-value less than 0.05 were defined as AZ-region associated. A greater number of AZ-region associated modules and notably more genes were observed within these modules in the *aus* lineage (six modules; Figure 3a-3b; 3e-3f and 3i-3j) compared to the *japonica* lineage (four modules; Figure 3c, 3g-3h and 3k) (Table S3-6, Figure 3). These findings are consistent with the greater similarity in AZ morphology observed between BHA and *aus* than between SWR and *temperate japonica* (Figure 1).

**Figure 3.**
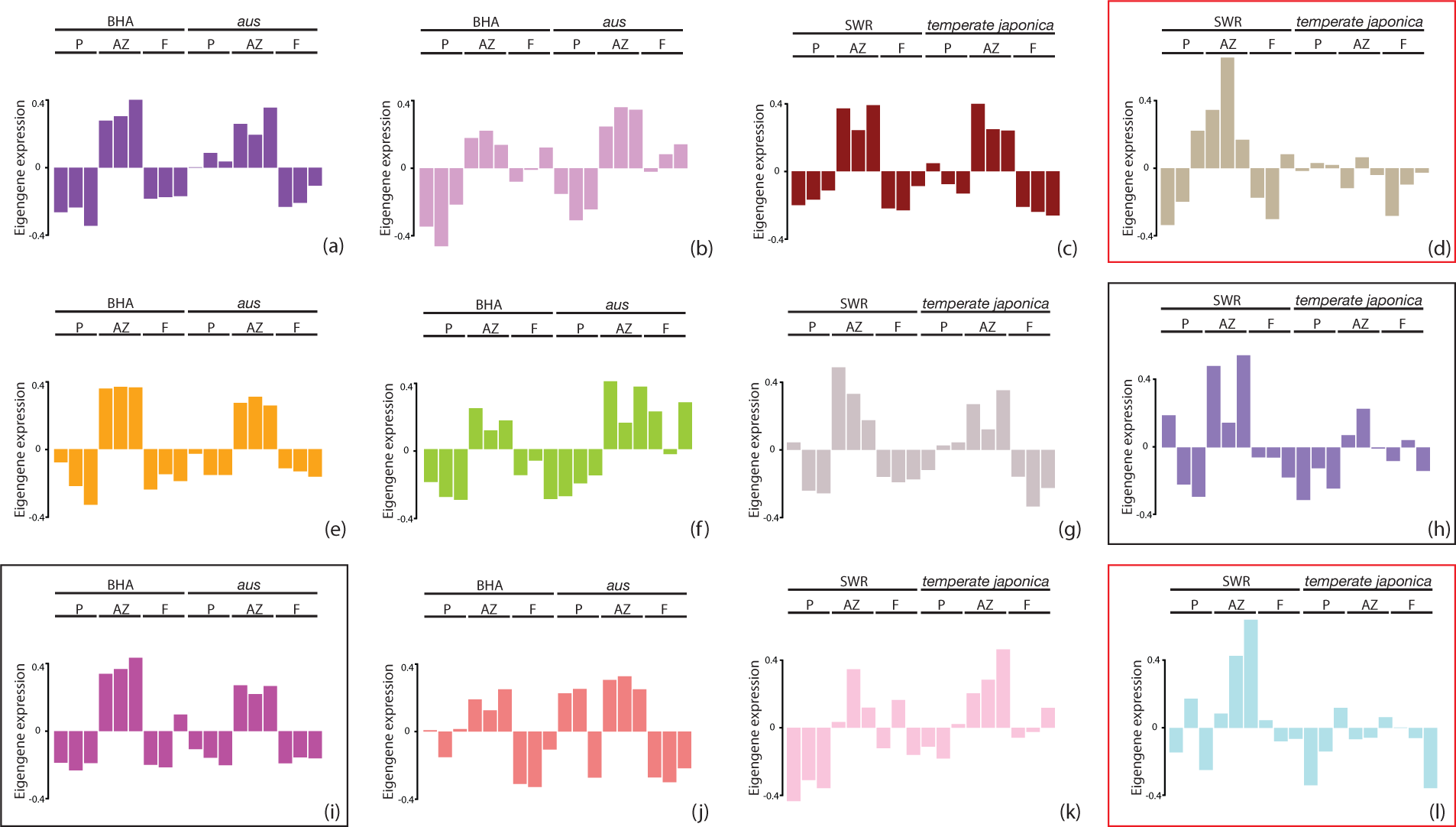
WGCNA modules associated with the AZ regions in both weedy and cultivated genotypes or in weedy genotype only. Eigengene expression boxplot for modules associated with AZ regions (AZ-region associated modules): (a-b, e-f, and i-j) modules associated with the AZ regions in BHA and *aus* genotypes detected at BFT, FT and LFT developmental stage respectively, a) purple module with 476 genes, b) plum module with 55 genes, e) orange module with 133 genes, g) yellowgreen module with 104 genes, i) magenta module with 215 genes and j) lightcoral module with 56 genes; (c-d, g-h and k-l) modules associated with the AZ regions in SWR and *temperate japonica* genotypes detected at BFT, FT and LFT developmental stage respectively, c) firebrick4 module with 74 genes, d) floralwhite module with 101 genes, g) lavenderblush3 module with 68 genes, h) mediumpurple2 module with 62 genes, k) lavenderblush1 module with 46 genes and l) powderblue module with 31 genes. The modules surrounded by red rectangle are associated with the AZ region in weedy genotype only. The modules surrounded by black rectangle are associated with the AZ region in both genotypes as well as having a distinct association with the AZ region in weedy genotype only. The remaining modules are modules associated with the AZ region in both genotypes.

We also identified modules exhibiting an AZ-region exclusive expression pattern specific to weedy rice but not to cultivated rice, through a separate correlation analysis assigning 1 to samples from the AZ region in weedy genotypes and 0 to all the other tissues in cultivated and weedy genotypes (Table S3). Genes in these modules may contribute to the development, maintenance, or competence of a complete AZ in weedy rice, and to weedy rice’s greater shattering ability; the altered expression patterns of these genes in cultivated rice may also be responsible for the morphological variation observed in the AZ region of cultivars. We identified weed-specific AZ-region associated modules with two types of expression patterns (Figure 3). One type comprised genes showing a distinctive AZ-region exclusive expression pattern solely in the weed, with no expression in the crop (Table S7; Figure 3d and 3l). The other type of module displayed the same AZ-region exclusive expression pattern in both the weed and crop, but with higher expression in the weed (Figure 3i and 3h); these modules were thus also detected in the correlation analyses involving crop and weedy genotypes concurrently. In the *japonica* lineage, three modules, one for each developmental stage, exhibited significant correlations with the weedy AZ region, with two being the first type module (Figure 3d and 3l) and one being the second type (Figure 3h). In the *aus* lineage, only one module detected in the later developmental stage, was significantly correlated with the weedy AZ region, exhibiting the expression pattern of the second type (Figure 3i). These results suggest that the same genes or genetic networks are involved in AZ development in BHA and *aus*, but it is their varying expression levels that lead to the slight differences in AZ morphology and the differences in seed shattering (Figures 1 and S1). Conversely, the restoration of a complete AZ in SWR seems to entail activation of genetic networks at multiple developmental stages that are largely not active in cultivated *temperate japonica*.

### The occurrence of genes related to lignin and cell wall modification within AZ-region associated modules

We examined each AZ-region associated module for enriched gene ontology (GO) terms, including modules associated in both cultivated and weedy genotypes and those associated with the AZ region only in weedy genotypes. Many modules did not have any significant GO-term enrichment and the number of enriched terms, when present, was usually modest, and primarily associated with various metabolic processes which did not provide meaningful insights into AZ-specific biological processes (Table S8 and S9, Figure S4 and S5). Given the observed characteristics of AZ-type cells, we then identified genes in each AZ-region associated module with GO terms related to lignin biosynthesis and cell wall modification. In total we found 8 and 24 genes associated with lignin metabolic process (GO:0009808) and cell wall organization and biogenesis (GO:0071554, GO:0042546 or GO:0009827), respectively (Table S10).

In the *aus* lineage, genes related to lignin modification were present in four out of six AZ-region associated modules for both genotypes, including the module more highly expressed in the BHA weedy genotype AZ region (Table S10). Surprisingly, in the *japonica* lineage, only two AZ-region associated modules for both genotypes identified in the BFT and FT stages contained lignin modification related genes, and none of the weed-specific AZ-region associated modules, despite the clear difference in lignified cells in the AZ region between cultivated and weedy rice (Table S10; Figure 1). Notably, *OsWRKY102*, a transcriptional repressor of secondary cell wall formation (Miyamoto *et al*., 2020), was identified in the AZ-region associated modules in BHA and *aus* genotypes during both the FT and LFT stages, and in SWR and *temperate japonica* genotypes during the FT stage. This gene was also present in the module associated with the AZ region only in BHA genotype (Table S10). A loss-of-function mutation in this gene results in increased lignin levels in rice (Miyamoto *et al*., 2020), thus the exclusive expression of *OsWRKY102* in the AZ region suggests its necessity in repressing lignin deposition. Its expression in the AZ region of the *temperate japonica* genotype, which does not form a distinct AZ cell layer but contains only a few AZ-type cells, suggests this gene is important for maintaining overall AZ region identity.

Cell wall modifications were regulated throughout all three developmental stages in both lineages. Generally, more genes involved in cell wall modification were found in AZ-region associated modules during the BFT stage compared to the later developmental stages, particularly in the BHA and *aus* genotypes (Table S10). During the BFT stage, the modification of pectin, regulating intercellular adhesion, appeared to be important for AZ development. *Pectin Methylesterase Inhibitor* (*PMEI*) genes, *OsPMEI 17* and *OsPME136*, appeared as hub genes, that is, genes with the highest top 10% connectivity within AZ-region associated modules, detected for BHA and *aus* genotypes and for SWR and *temperate japonica* genotypes during this stage (Table S6 and S10). Several genes related to cell wall modification were also found within the weed-exclusive AZ-region associated modules (Table S10). For instance, LOC_Os02g32110 in the SWR AZ-region associated module is a homolog of *AtIRX10* in *Arabidopsis thaliana*, which regulates the synthesis of xylan, a hemicellulose in the cell wall, and exhibits a conserved function over a substantial evolutionary distance (Jensen *et al*., 2014). Another example was a member of the xyloglucan endotransglucosylase/hydrolase (*XTH*) gene family, *XTH3*, in the BHA AZ-region associated module. Genes in the *XTH* family are involved in the biosynthesis of xyloglucan, another type of hemicellulose in cell wall (Yokoyama *et al*., 2004).

### Genes overlapping in the AZ-region associated modules among different rice lineages

There was limited overlap of genes within AZ-region associated modules between the *aus* and *japonica* lineages (Table S11). The highest number of overlapping genes, 14, was observed at the BFT stage (Figure 4, Table S11). Notably, an APETALA2-like transcription factor (LOC_Os03g60430) exhibiting a high degree of similarity to known seed shattering genes, *SHAT1* and *SNB*, which are involved in AZ formation (Wang *et al*., 2019), was detected in AZ-region associated modules in both lineages (Table S11). This gene also served as a hub gene in the AZ-region associated module for BHA and *aus* genotypes during this BFT stage (Table S6). The pectin regulation gene *OsPMEI36* also overlapped between modules of the two lineages during the BFT stage (Table S11). These results suggest that at this stage, determining AZ region identity and modifying cell wall composition are likely shared processes between these two lineages.

**Figure 4.**
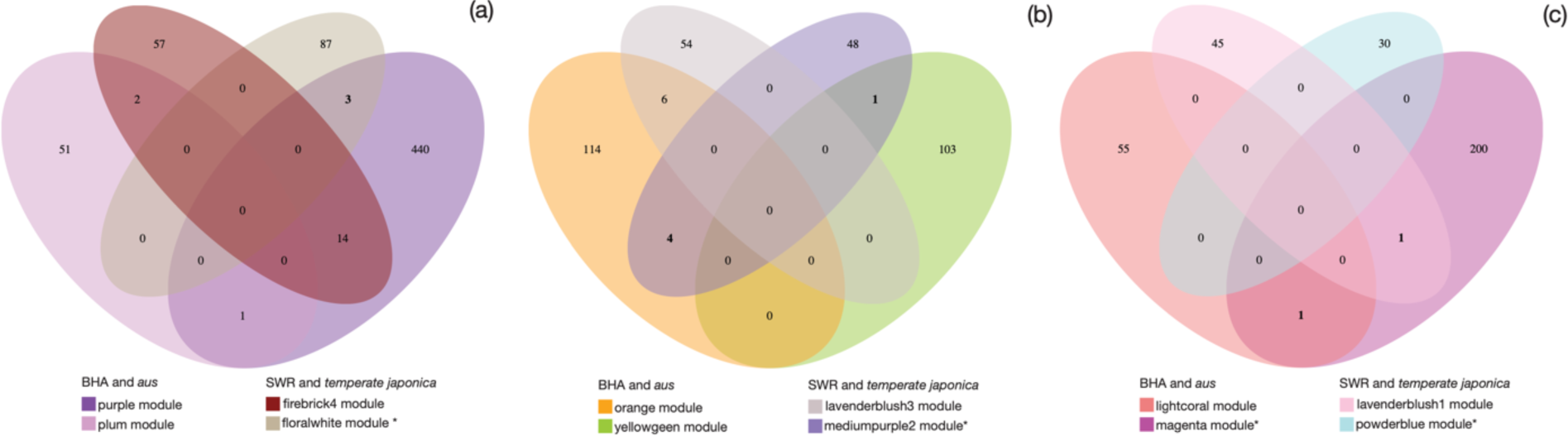
The number of overlapping genes within the AZ-region associated modules between *aus* and *japonica* lineages at each developmental stage. (a) overlap at the BFT stage; (b) overlap at the FT stage; and (c) overlap at the LFT stage. Ellipses representing AZ-region associated modules detected in SWR and *temperate japonica* are positioned inside while ellipses representing AZ-region associated modules detected in BHA and *aus* are positioned outside. Legends for modules associated with the AZ region in weedy genotype only are denoted with “*”. The number of genes overlapping between modules associated with the AZ region in the weedy genotype only and modules associated with the AZ region in both weedy and cultivated genotypes are bolded.

At the FT stage, three genes stood out among those overlapping between the AZ-region associated modules in both lineages: *WRKY102*, *GA2-oxidase7* (*GA2ox7*) and a NAC transcription factor (*OsXND1-like*) (Table S11). *WRKY102* functions as lignin biosynthesis repressor (Miyamoto *et al*., 2020). *GA2ox7* codes for an enzyme involved in regulating gibberellin (GA), and modulating GA content can impact lignin deposition and ultimately alter seed shattering (Wu *et al*., 2023). *OsXND1-like* is homologous to *A. thaliana* Xylem NAC domain 1 (*XND1*), which negatively regulates secondary cell wall synthesis and promotes programmed cell death in xylem (Zhao *et al*., 2008). Ectopic expression of *XND1* in fiber cells failed secondary cell wall deposition and dramatically reduced the expression of secondary wall biosynthesis genes (Zhao *et al*., 2008). This gene was hub gene in the AZ-associated modules detected in both the BHA and *aus* genotypes as well as SWR and *temperate japonica* genotypes at this developmental stage (Table S6). Importantly, this gene ranked as the second most interconnected gene within the AZ-associated module detected in the BHA and *aus* genotypes (Table S6). These findings indicate that the primary processes shared among these four genotypes during the FT developmental stage included the repression of lignin deposition and the inhibition of secondary cell wall formation in the AZ region.

During the LFT stage, only two genes, specifically a protein kinase and a membrane protein, exhibited overlap (Table S11). Compared to the other developmental stages, there was a reduced number of shared genes among these genotypes at this stage.

### Genes overlapping across development in AZ-region associated modules

There was no overlap observed among AZ-region associated modules across all three developmental stages, regardless of whether they were exclusively associated with the weedy AZ region or simultaneously associated with both weedy and cultivated AZ regions (Figure 5). More genes were shared between AZ-region associated modules from different developmental stages in the *aus* lineage compared to the *japonica* lineage (Table S12).

**Figure 5.**
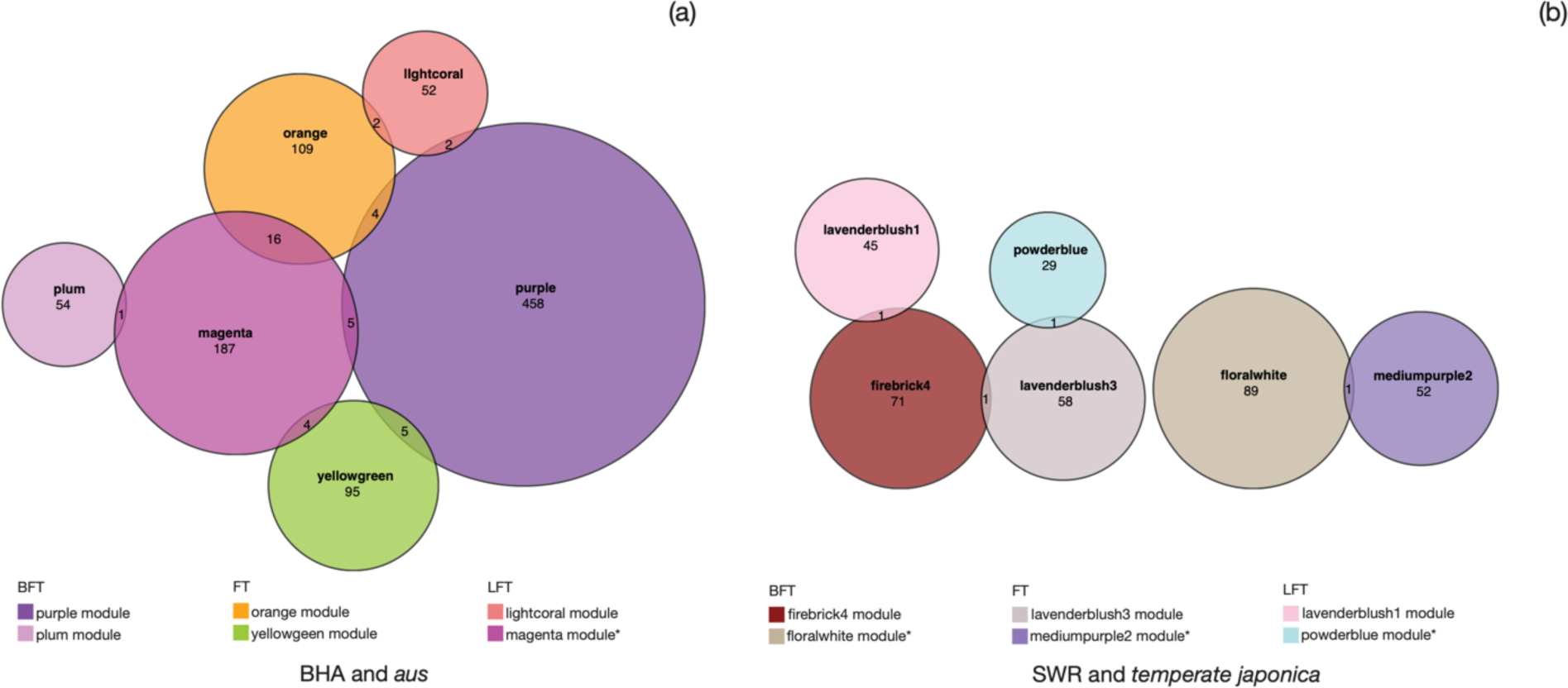
The number of overlapping genes among the AZ-region associated modules detected within each lineage across development. (a) overlap of AZ-region associated modules detected within the *aus* lineage; and (b) overlap of AZ-region associated modules detected within the *japonica* lineage. Legends for modules associated with the AZ region in weedy genotype only are denoted with “*”.

The most substantial overlap (16 genes) was observed between the FT (orange) and LFT (magenta) AZ-region associated modules in the *aus* and BHA genotypes (Figure 5), in line with the results of the PCA where samples from these two stages clustered together (Figure 2a). Interestingly, the genes that overlapped between these two AZ-region associated modules included four out of six genes that also overlapped between AZ-region associated modules from different lineages during the FT developmental stage (Table 12). This included one coding for a MATE efflux family protein and three previously mentioned genes, *WRY102*, *GA2ox7* and *OsXND1-like*, suggesting a consistent requirement for repressing lignin deposition and secondary cell wall deposition and the potential involvement of the same genetic networks in regulating these processes in the AZ in both developmental stages in BHA and *aus*. However, only one gene, *ubiquitin domain-containing protein 2*, overlapped between FT and LFT stages in the AZ-region associated modules identified in the *japonica* lineage (Table 12).

### Previously known seed shattering genes exhibiting tissue-specific expression patterns

Several genes associated with seed shattering levels in rice have been previously identified via crosses or mutagenesis (Turra *et al*., 2023). We examined modules containing these previously identified genes to determine how they related to the shattering differences observed among crops and weeds in each of our lineages. We particularly focused on the genes that have been functionally verified using transformation. Four genes - *SH4* (Li *et al*., 2006), *qSH1* (Konishi *et al*., 2006), *OsSh1* (Li *et al*., 2020) and *OsLG1* (Ishii *et al*., 2013) - were primarily identified through crosses, and natural variation within these genes is associated with reduced seed shattering in cultivated rice during domestication. Eight genes, including *SH5* (Yoon *et al*., 2014), *SNB* (Jiang *et al*., 2019), *SHAT1* (Zhou *et al*., 2012), *OsCPL1* (Ji *et al*., 2010), *OSH15* (Yoon *et al*., 2017), *SLR1* (Wu *et al*., 2023), *OsGRF4* (Sun *et al*., 2016) and *NPC1* (Cao *et al*., 2016), were mainly identified through mutagenesis, with their roles in domestication remaining unknown or unexamined.

We found that most of these known shattering candidate genes occurred in modules that were expressed in all three developmental stages, with the exception of *SH4* and *OsLG1* (Figure S6). The eigengenes of the majority of these modules containing candidate genes exhibited a tissue-specific expression pattern, with non-uniform expression across all three tissues, (Figure S6), suggesting that the specific location of gene expression was crucial for fulfilling their functions. The interactions among these genes were also reflected in their memberships within the same module. For example, *OsSh1*, *SNB* and *qSH1* were included in the same module detected in BHA and *aus* during BFT (Figure S6). While *OsSh1* and *qSH1* were in the same module detected in SWR and *temperate japonica* during BFT; *OsSh1* and *SNB* were in the same module detected in SWR and *temperate japonica* during LFT (Figure S6). These results suggested that although these genes work together in both lineages, the nature of interactions might differ.

Because most of these genes have been previously implicated in AZ formation in early floral development, we focused on their expression patterns during the BFT stage. *qSH1* (Konishi *et al*., 2006) and *OsSh1* (Li *et al*., 2020; Ishikawa *et al*., 2022) were detected in modules demonstrating high expression in the AZ and floret tissues in the BHA, *aus* and SWR genotypes, while no expression in the AZ was observed in the *temperate japonica* (Figure 6a). A similar situation where high expression in the AZ and pedicel tissues was evident in the rice varieties with a well-defined AZ morphology, BHA, *aus*, and SWR, while no AZ expression was discernible in the *temperate japonica*, was also observed for the modules that contained *SH5* (Yoon *et al*., 2014) (Figure 6b) and *NPC1* (Cao *et al*., 2016) (Figure 6c). The *qSH1* and *OsSh1* containing modules (Figure 6a) overlapped by 258 genes between the two lineages and the *NPC1* containing modules (Figure 6c) overlapped by 158 genes. These findings suggest that the genes within these modules might form certain genetic networks and be crucial for an AZ layer presence and shattering in both lineages.

**Figure 6.**
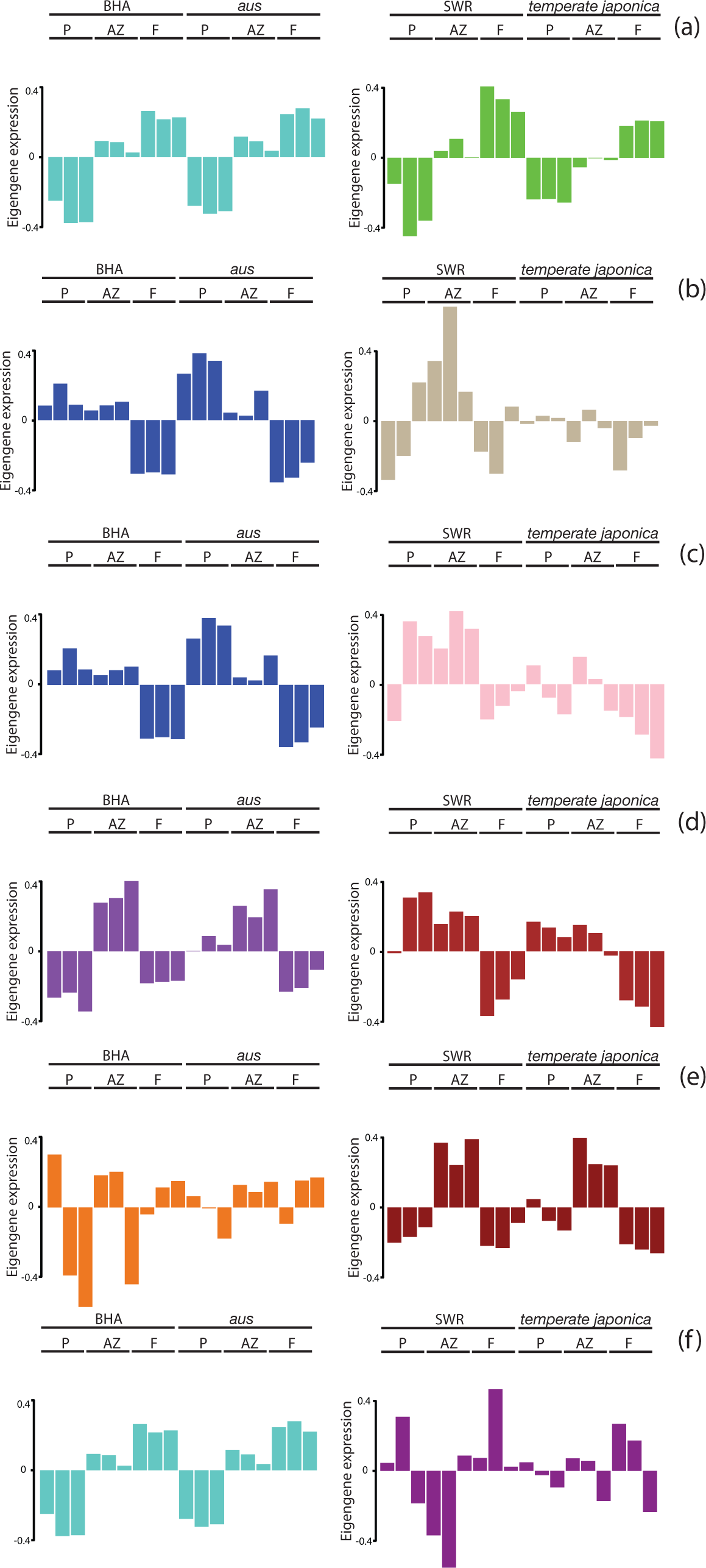
WGCNA modules at the BFT stage containing known seed shattering genes. Eigengene expression boxplot for modules including the known seed shattering genes: (a) modules including *qSH1* and *OsSh1* detected in the *aus* lineage and the *japonica* lineage respectively at BFT stage; (b) modules including *SH5* detected in the *aus* lineage and the *japonica* lineage respectively at BFT stage; (c) modules including *NPC1* detected in the *aus* lineage and the *japonica* lineage respectively at BFT stage; (d) modules including *SHAT1* and *OSH15* detected in the *aus* lineage and the *japonica* lineage respectively at BFT stage; (e) modules including *SH4* detected in the *aus* lineage and the *japonica* lineage respectively at BFT stage; and (f) modules including *SNB* detected in the *aus* lineage and the *japonica* lineage respectively at BFT stage. Grey are the modules where unassigned genes were labeled.

We also found that certain AZ genes were part of modules with a consistent expression pattern between the crop and weed with the same genetic background but exhibited shifted expression patterns in rice with different genetic backgrounds. For example, *SHAT1* (Zhou *et al*., 2012) and *OSH15* (Yoon *et al*., 2017) were both members of a module exclusively expressed in the AZ in the BHA and *aus* genotypes, but were detected in a module expressed in both AZ and pedicel tissues in the SWR and *temperate japonica* genotypes (Figure 6d). 35 genes were overlapped between these *SHAT1* and *OSH15* containing modules (Figure 6d) between the two lineages. Conversely, *SH4* (Li *et al*., 2006) occurred in a module displaying exclusive expression in the AZ in the SWR and *temperate japonica* genotypes, while the *SH4* containing module in the BHA and *aus* genotypes did not exhibit a clear pattern, possibly showing higher expression in both AZ and floret tissues (Figure 6e). There were also situations where a gene belonging to a module demonstrating high expression in the AZ and floret tissues in one lineage, displayed a relatively random expression pattern in the other lineage; *SNB* (Jiang *et al*., 2019) was one such example (Figure 6f). Few genes of these modules (Figure 6e - 6f) overlapped between the two lineages.

### Genes exhibiting differential expression in the AZ between weedy and cultivated rice in the ***aus* and *japonica* lineages**

To identify genes potentially responsible for variation in seed shattering between cultivated and weedy rice within each lineage, we performed differential expression analyses for each tissue type at every developmental stage. Hundreds of genes were differentially expressed (DE) between crops and weeds in every tissue and at every stage. We focused on DE genes within the AZ and found larger number between BHA and *aus* -- 545, 712 and 642 at the BFT, FT and LFT developmental stages respectively (Table S14) -- than between SWR and *temperate japonica* -- 420, 312 and 368 respectively (Table S15). None of known shattering genes exhibited significant expression difference within the AZ between crop and weed in any lineage. This difference in DE gene number, however, was reduced once we examined genes exclusively DE in the AZ but not in the other tissues; 132, 234, and 133 AZ-exclusive DE genes were identified between BHA and *aus,* while 162, 70 and 152 were found between SWR and *temperate japonica* at the BFT, FT and LFT developmental stages, respectively (Table S16). Among these AZ exclusive DE genes, 28, 30, and 26 genes overlapped at each stage between the weed-crop comparisons, although the directions of regulation of some of these genes were not consistent between comparisons (Table S17).

### GO term enrichment for weedy-cultivated rice DE genes

Several related GO terms were consistently enriched for AZ-exclusive DE genes between BHA and *aus* across all three developmental stages, such as “cell death”, “programmed cell death”, “apoptosis” and “response to stress” (Figure 7 and Table S18). Numerous GO terms related to metabolic and cellular regulation were exclusively enriched during the FT stage (Figure 7 and Table S18). In contrast, no enriched GO terms for the AZ-exclusive DE genes between SWR and *temperate japonica* were shared across the three developmental stages. During the FT stage, only a single GO term, “response to biotic stimulus”, displayed enrichment. During the BFT stage, multiple GO terms related to “photosynthesis” and one GO term related to “cell-wall” were enriched. During the LFT stage, the enriched GO terms encompassed “response to stress/stimulus”, “DNA packaging” and “DNA conformation change” (Figure 7 and Table S18). Cell death related GO terms were not enriched for AZ-exclusive DE genes between SWR and *temperate japonica*. The only shared enriched GO terms with the BHA-*aus* comparison were “response to stress” and “response to stimulus.” This suggests that different regulatory processes are responsible for the differences in seed shattering and AZ morphology between crops and weeds in each rice lineage. Particularly, the enriched GO terms in the SWR and *temperate japonica* comparison appeared to be more focused on responses while those in the BHA and *aus* comparison were more related with cell identity (Figure 7).

**Figure 7.**
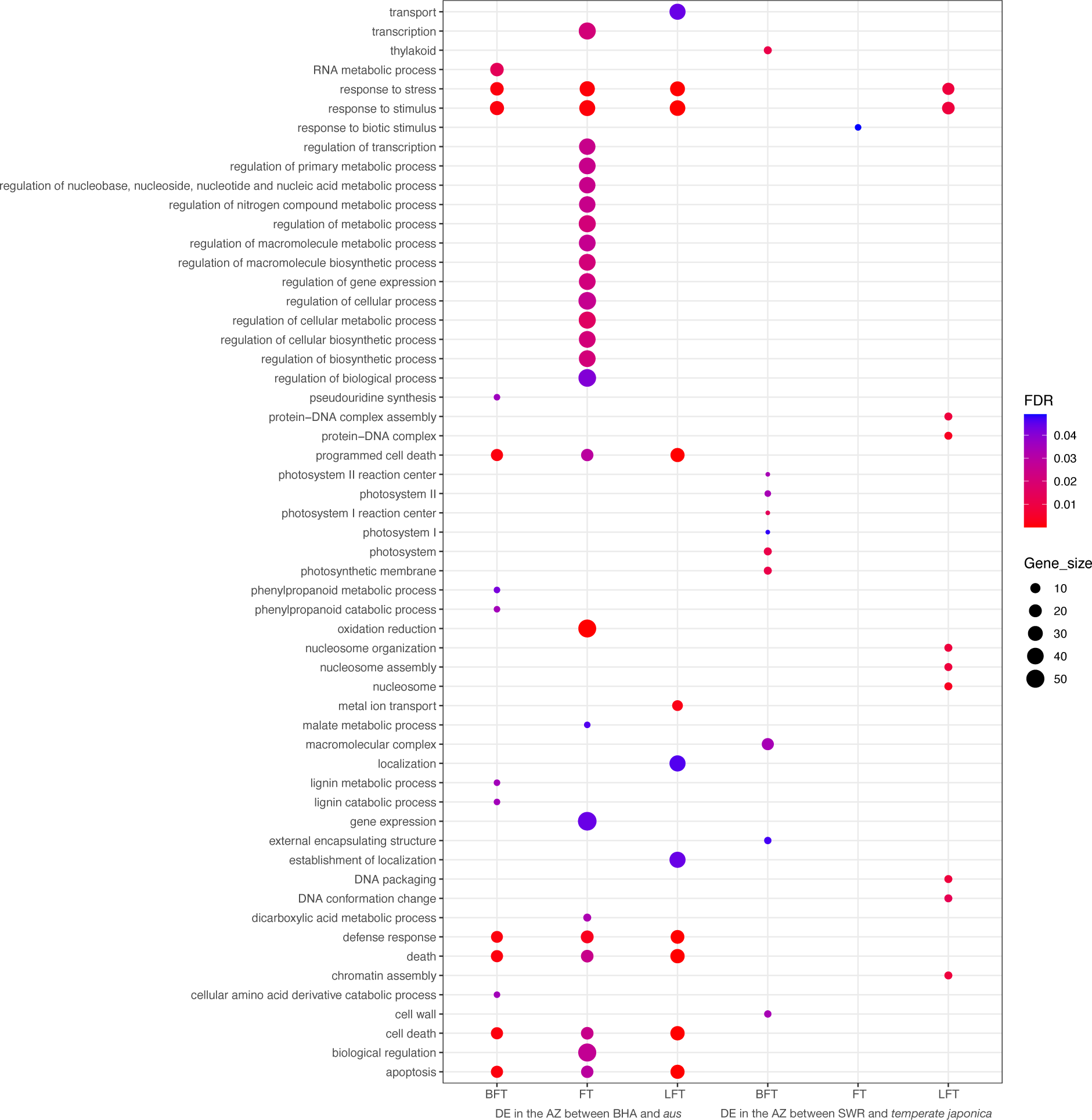
Functional categories enriched in AZ region differentially expressed (DE) genes between each weedy-cultivated comparison at each developmental stage. Significant Gene Ontology (GO) terms enriched in the DE genes in the AZ between BHA and *aus* as well as SWR and *temperate japonica* at BFT, FT and LFT developmental stages. Red and blue represent smaller and larger FDR values, respectively. The size of the dots represents the number of genes in each GO term over the total number of DE genes in the AZ.

### Genes related to lignin modification differentially regulated between weedy and cultivate rice

Similar to the GO analysis in AZ-region associated modules, we examined genes annotated with GO terms of lignin metabolic process (GO:0009808) and cell wall organization or biogenesis (GO:0071554) among the DE genes exclusively in the AZ region in each weedy-cultivated comparison (Table S18).

During the BFT stage, three laccase genes (*OsLACs*) exhibited downregulation in the AZ region of BHA compared to *aus* (Table S19). Laccases, a type of oxidative enzyme, play a crucial role in determining the spatial and temporal polymerization of lignin (Emonet and Hay, 2022). The diminished activity of these genes may lead to less lignin being deposited in the AZ of BHA, thereby contributing to its increased shattering compared to *aus*. Notably, *OsLAC15_2* consistently demonstrated reduced expression in the AZ of BHA compared to *aus* across all three developmental stages, suggesting a persistent reduction in lignin deposition in the AZ region of BHA throughout development. In line with this lignin repression, *OsWRKY102*, a repressor of lignin deposition, was found to be highly expressed in the AZ of BHA compared to *aus* during the FT stage (Table S19). In contrast, there were fewer DE genes in the AZ region between SWR and *temperate japonica* associated with lignin (Table S19). One of these genes was *OsWRKY36*, another repressor for lignin and secondary cell wall deposition (Table S19). *OsWRKY36* activates the transcription of *SLR1* by directly binding to its promoter (Lan *et al*., 2020); *SLR1*, in turn, functions as a key repressor of GA signaling directly involved in seed shattering in rice through impacting lignin content in the AZ region (Wu *et al*., 2023). Interestingly, *OsWRKY36*, displayed lower expression in the AZ of SWR compared to *temperate japonica* during the LFT developmental stage (Table S19), despite lower levels of lignin expected in SWR.

*Pectin methylesterase* (*PME*), *XTH* genes, *cellulose Synthase - like* (*CSL*) genes, involved in biosynthesis of pectin, hemicellulose, and cellulose respectively, exhibited differential gene regulation between weedy and cultivated rice in both lineages (Table S19). The modification of these components can impact the deposition of secondary cell wall (Coomey *et al*., 2020). However, examining the precise impact of these genes on cell wall modifications can be challenging, given that the function and interactions of these genes have not be well-characterized in rice.

## Discussion

Seed shattering is one of the most consistently differentiated traits that distinguishes weedy from cultivated rice, with high seed shattering significantly increasing the competitiveness of weeds in rice fields (Vigueira *et al*., 2013). The repeated evolution of high shattering in de-domesticated weedy rice stemming from different cultivated ancestors provides an excellent framework for exploring mechanisms of convergent evolution. Our previous examinations of AZ histology suggest that AZ-type cells converge in their cellular characteristics, irrespective of genetic background; in contrast, the arrangement of AZ-type cells, in particular the length of the AZ cell layer, is associated with different levels of seed shattering across different weedy and cultivated rice populations (Li *et al*., 2024). Examining patterns of gene expression in the AZ region of related weedy-cultivated rice pairs enabled us to identify the genetic associations with AZ morphology and shattering levels in the *aus* and *japonica* lineages, and the extent of genetic convergence. Our findings indicate constraints on the genetic mechanisms determining AZ-type cell identity, but flexibility in the genetic networks shaping AZ morphology, particularly concerning AZ-type cell distribution, underscoring heterogeneity in the levels of genetic convergence underlying the crucial agricultural trait of shattering.

### Genetic convergence underlying the maintenance of AZ identity across lineages

AZ-type cells exhibit uniform cellular characteristics, featuring the absence of lignin, small size and thin walls, across our four examined rice genotypes (Figure 1). These characteristics appear very early and persist throughout floral development (Figure 1), emphasizing the necessity in maintaining AZ-type cell identity. Some genes related to lignin and secondary cell walls, especially transcription factors, consistently display comparable AZ-exclusive or tissue-specific expression patterns in both weedy and cultivated genotypes across lineages. These genes are likely instrumental in determining the distinct AZ-type cell identity, as evidence of genetic convergence in both lineages.

One such example is a gene encoding an *AP2*-like transcription factor detected in the AZ-region associated module for weedy and cultivated genotypes in both lineage during the BFT stage (Table S11). This gene exhibits high similarity to *SHAT1* and *SNB,* both involved in AZ formation, the latter of which can inhibit lignin deposition in the AZ (Jiang *et al*., 2019). Another example is *OsWRKY102*, a repressor for lignin biosynthesis, which not only appears in AZ-region associated modules from both lineages during the FT stage (Figure 4, Table S11), but also preserves this AZ-exclusive expression pattern in BHA and *aus* into the LFT stage (Figure 5, Table S12). Additionally, a higher expression of this gene is observed in the AZ of BHA compared to *aus* (Table S19), correlated with the greater number of AZ-type cells in BHA, an expected outcome of more repression of lignin deposition. These findings suggest that lignin deposition remains regulated consistently throughout AZ development. Nevertheless, this repression may undergo some changes, as evidenced by the observed decrease of expression of *OsWRKY102* during the LFT stage compared to the FT stage in BHA and *aus*. Repression of secondary cell wall deposition is another process consistently regulated across AZ development. One transcription factor, *OsXND1-like*, associated with repressing secondary cell wall deposition, occurred in the AZ-region associated modules in both lineages during the FT stage (Table S11) and in BHA and *aus* at the LFT stage (Table S12). Several cell wall modification enzymes also exhibited AZ-exclusive expression pattern in both lineages (Table S10) supporting genetic convergence.

Certain known seed shattering genes interact, and the outcomes of their interactions include the alteration of lignin or secondary cell wall deposition. Although these genes did not exclusively express in the AZ region, they exhibited a tissue-specific expression pattern and possibly interacted with each other if they were in the same module. For example, *OsSh1* functions downstream of *qSH1* (Li *et al*., 2020) while *qSH1* is directly regulated by *SNB* (Jiang *et al*., 2019). The direct impact of their interaction is the suppression of lignin deposition in the AZ region, and mutations in *SNB* lead to altered development of AZ (Jiang *et al*., 2019). These three genes were found in the same module, exhibiting higher expression in the AZ region and floret tissues in BHA and *aus* during the BFT stage (Figure S6); additionally, *qSH1* and *OsSh1* are in the same module in SWR and *temperate japonica* during BFT, and *OsSh1* and *SNB* are in the same module in these two genotypes during LFT (Figure S6). *NPC1* can impact the secondary cell wall deposition through modulating silicon distribution and resulting in changes in the content of cellulose and hemicellulose in the cell wall and the thickness of cell wall (Cao *et al*., 2016). This gene is included in the module exhibiting higher expression in the pedicel and AZ region during the BFT stage in BHA, *aus* and SWR (Figure 6c), and potentially interacts with *SH5* in BHA and *aus* (Figure 6b). Additionally, relatively large number of genes overlapped between the *qSH1* and *OsSh1-*containing and *NPC1-*containing modules from the two lineages (Figure 6a and 6c).

Although indirect, these results suggest that a shared group of genes, particularly transcription factors, are involved in determining AZ region identity in different genotypes of rice, indicating possible genetic constraints to making AZ-type cells in rice.

### Genetic divergence underlying AZ variation between weedy and cultivated rice in different lineages

How the AZ morphology varies between cultivated and weedy rice is lineage-dependent, suggesting that distinct genetic mechanisms explain these weed-crop differences. The primary AZ morphological distinction between BHA and *aus* lies in the length of the AZ (Figure 1), indicating that the genetic basis for AZ region formation is likely similar, with differences possibly in allowing or repressing AZ formation close to the vascular bundle (Figure 1g-1i). This aligns with our finding that all AZ-region associated modules were consistently correlated with the AZ region in both BHA and *aus* (Figure 3). Additionally, the modules including previously known seed shattering genes largely exhibited similar tissue-specific expression patterns between these two genotypes (Figure S6). Thus, the same networks appear to be utilized in both genotypes to maintain AZ cell identity, but these genes might exhibit varied expression levels and result in the AZ morphological difference and seed shattering difference between them. One possible causal gene might be *SHAT1*, a master transcription factor determining AZ identity (Zhou *et al*., 2012). Although not significant, this gene exhibits higher expression in the AZ in BHA than in *aus*. This gene is also in a seed shattering QTL detected through a cross of the same BHA and *aus* genotypes from this study and this QTL overlaps with a selective sweep (Li *et al*., 2023). One single nucleotide mutation in the 3’ UTR (untranslated region) of this gene exhibited association with the change of seed shattering levels (Li *et al*., 2023).

In contrast to weeds and crops in the *aus* lineage, substantial morphological distinctions are present in the AZ region between SWR and *temperate japonica*. The discontinuous AZ in *temperate japonica* (Figure 1) suggests that AZ-type cells are randomly scattered in the AZ region, lacking the well-defined AZ cell layer observed in *aus*. Consistent with this disrupted or underdeveloped AZ region in *temperate japonica*, fewer AZ-region associated modules were shared by both genotypes in the *japonica* lineage compared to the *aus* lineage. Additionally, modules with AZ-region exclusive expression in weeds but no expression in crop were only detected in the *japonica* lineage (Figure 3). The random and scattered distribution of AZ cells in *temperate japonica* suggests inconsistent regulation of AZ cell formation in this genotype. For example, the module that included *OsSh1* and *qSH1* in the BFT stage, displayed comparable tissue-specific expression pattern in BHA, SWR and *aus*, while an altered expression pattern in *temperate japonica* (Figure 6a).

That different genes have been involved in the re-evolution of easy shattering in different de-domestication events of weedy rice had also been previously suggested by the lack of overlapping shattering QTLs detected in mapping population from weed x crop crosses involving BHA weedy rice, and SH weedy rice - a group descended from *indica* domesticated varieties (Thurber *et al*., 2013; Qi *et al*., 2015; Li *et al*., 2023). Different sets of genes exhibiting differential expression between crop and weed within each lineage further illustrate that different genes involved in the reacquisition of high shattering in these two independently evolved weedy rice populations.

Overall, our results shed light on the genetic convergence underling phenotypic convergence: consistent repression of lignin and secondary cell wall deposition throughout development is crucial for maintaining the identity of the AZ region, regardless of weedy or cultivated rice genotype; this phenotypic convergence is supported by the observation that some genes associated with these modifications exhibit exclusive expression in the AZ region across both weedy and cultivated genotypes. However, only a few of these genes overlap between the two lineages, suggesting that the specific genes responsible for modifying lignin and secondary cell walls may differ between them. This variation could be attributed to distinct genetic backgrounds or evolutionary histories.

## Experimental Procedures

### Plant materials and sample collection

Two cultivated rice lines, Aus 196 and AN-09-13 were chosen to represent the rice variety groups of *aus* and *temperate japonica*. Lines 10A and ASF-09-54, were chosen to represent the two weedy rice populations Black Hull Awned (BHA) and Spanish Weedy Rice (SWR), which evolved from *aus* and *temperate japonica*, respectively. Aus 196 is a breeding and research material developed in Bangladesh and acquired from the International Rice Research Institute; 10A is a weedy rice variety collected in Arkansas, U.S. and maintained by the United States Department of Agriculture; AN-09-13 and ASF-09-54 were obtained through field collections and sampling of germplasm repositories in the Iberian Peninsula (Li *et al*., 2022). At least two seeds of the same genotype were planted in one pot; each pot served as one replicate for each genotype; and three replicates were used for each genotype in the RNA-seq experiments. The plants were grown under their optimal growth conditions (11 hour light and 12 hour dark with the day temperature of 29 ℃ and the night temperature of 24℃) in a growth chamber at the University of Massachusetts Amherst.

### Histology and seed shattering measurements

Histological analyses were conducted following established procedures as described in (Li *et al*., 2024). Briefly, florets were collected at three distinct developmental stages for each genotype: one week before the heading date (BFT), the heading date (FT) and one week after the heading date (LFT). Heading date refers the stage when the first panicle has emerged halfway from the boot. These florets were fixed in an FAA solution, dehydrated in a graduated ethanol series, cleared with Histoclear, embedded in paraplast, and eventually sectioned into 10 *μm* longitudinal slices with a Leica RM2125 microtome. The resulting sections were stained with 1% toluidine blue and observed using a Leica DM750 LED biological microscope. High-quality images were captured using an AmScopre MU1000-HS camera.

Seed shattering levels were evaluated by measuring the breaking tensile strength (BTS), the force required to separate the seeds from the plants. This measurement was performed 30 days after heading (LFT30), a point at which the BTS values stabilize (Thurber *et al*., 2010; Thurber *et al*., 2011). For each panicle, 10 florets were assessed and at least two panicles per genotype were included in the measurement. In addition to measuring BTS at LFT30, we also conducted BTS measurements at FT and LFT to capture the trends in BTS changes during development. Notably, during the BFT stage, the spikelet is exceptionally fragile, preventing the measure of BTS from this stage.

### RNA isolation, RNA-Seq library construction and sequencing

Florets were harvested from the four rice genotypes at the three developmental stages —BFT, FT and LFT— during the afternoons between 2:00 and 4:00 PM. For each developmental stage, we collected three replicates of three different tissues: AZ, the tissue toward floret side (F), and the tissue toward pedicel side (P). In each replicate, we combined tissues dissected from around 40-60 florets, typically sourced from at least two different plants grown in the same pot. The manual tissue dissection was carried out in RNAlater stabilization solution (Invitrogen) using a scalpel under a dissecting microscope (Leica), following the protocol described in (Yu *et al*., 2020). The dissected tissues were promptly transferred into RNAlater stabilization solution, stored at 4℃ overnight, and subsequently preserved at –80℃ before extraction. Dissected tissues were manually ground into powder using pellet pestles in liquid nitrogen. We performed RNA extraction using the PicoPure RNA Isolation Kit (Thermo Fisher Scientific) and included in-column DNase I (QIAGEN) treatment. The concentration and integrity of the RNA samples were assessed using the Nanodrop2000c (Thermo Fisher) and the Fragment Analyzer System with the DNF-471 RNA kit (15nt) (Agilent), respectively. For library preparation, we followed the manufacturer’s instructions for the QuantSeq 3’ mRNA-Seq FWD Library Prep Kit (Lexogen) (Moll *et al*., 2014). A total of 108 libraries (4 genotypes x 3 tissues x 3 developmental stages x 3 replicates) were single-end sequenced on a NextSeq 500 instrument at Lexogen (Vienna, Austria). The raw sequencing data will be deposited in the Sequence Read Archive.

### Reads alignment and principal component analyses

The raw reads for the four rice genotypes were cleaned with Trimmomatic using a sliding window trimming method with an average base quality threshold set at a minimum of 20 (Bolger *et al*., 2014). The resulting clean reads from the *aus* and its derived weedy rice BHA were mapped to the *indica* Shuhui 498 (R498) reference genome (Du *et al*., 2017) using Hisat2 (Pertea *et al*., 2016). The mapped reads were subsequently used as input for HTSeq-count (Anders *et al*., 2015) to calculate gene counts. The clean reads from the *japonica* and its derived weedy rice SWR were mapped to the Nipponbare *japonica* reference genome (Kawahara *et al*., 2013), following the same method as described above, to obtain gene counts for subsequent analyses.

Additionally, we also mapped the clean reads from the *japonica* and its derived weedy rice SWR to the R498 reference genome and conducted Principal Component Analysis (PCA) using the gene counts mapped to the same reference genome for all four genotypes to gain a comprehensive understanding of the expression pattern across the four genotypes.

### Weighted Gene Co-expression Network Analyses

We performed Weighted Gene Co-expression Network Analyses (WGCNA) for each cultivar and its corresponding derived weed pair (BHA and *aus*; SWR and *temperate japonica*) using all three tissues at each developmental stage to detect modules. Following the application of a variance-stabilizing transformation to the gene counts using the vst function in DESeq2 (Love *et al*., 2014), genes exhibiting low variance (Interquartile range < 0.5) were filtered out using the VarFilter function in the geneFilter package (Gentleman *et al*., 2023). The resulting gene counts were used for sample clustering and the construction of gene co-expression networks using the WGCNA package (Langfelder and Horvath, 2008). We employed the pickSoftThreshold function to choose the optimal power for the network and blockwiseModules function to construct a signed network with a mergeCutHeight of 0.25, a corType of “bicor” with maxPOutliers=0.05 and robustY=FALSE, and a minModuleSize of 30. The Pearson correlation coefficient between each module and AZ region in both genotypes or in the weedy genotype exclusively, as well as the associated statistical significance, were calculated using the cor and corPvalueStudent functions. Intramoduler connectivity and signed eigengene-based connectivity, also known as module membership, for each gene within their respective modules were determined using intramodular and signedKME functions.

### Differential expression analyses

We performed differential gene expression analyses between cultivated rice and its corresponding weedy rice in each tissue at each developmental stage using DESeq2 (Love *et al*., 2014). Genes with an average gene count greater than 10 in any specific tissue type at any developmental stage and in either genotype, cultivated or weedy, were included in the analyses. We applied a minimum two-fold change and an adjusted p-value below 0.05 as thresholds to determine genes with significantly differentiated expression.

### Gene function and GO enrichment analyses

Gene Ontology (GO) enrichment analyses were performed using the genes within each AZ-enriched module and the genes exhibiting differential expression in the AZ between cultivated rice and its corresponding derived weedy rice within each lineage using agriGO v2.0 with the setting of Hypergeometric as statistical test method, Hochberg (FDR) as multi-test adjustment method, 0.05 as significance level and one as minimum number of mapping entries (Tian *et al*., 2017). We referred to GO terms sourced from PLAZA 5.0 (Van Bel *et al*., 2022) to elucidate the functions of these genes.

## Acknowledgements

We thank the Caicedo Lab members Sherin Perera and Daniel Lowey for taking care of the plants; Dr. Elizabeth Kellogg and Dr. Yunqing Yu at Donald Danforth Plant Science Center for technical help; and Dr. Maria Dolores Osuna for providing seeds of Spanish weedy and cultivated rice. This work was supported by the National Science Foundation (NSF) [grant IOS-1947609 to A. Caicedo, K. Olsen, and Y. Jia], Lotta Crabtree Fellowships in Production Agriculture to XL, Gilgut Scholarships to XL, and UMass Biology Graduate Research Grant to XL.

## Supplementary Data

Table S1. The Breaking Tensile Strength (BTS) measurements for rice varieties across different developmental stages.

Table S2. PCA results for each accession.

Table S3. The AZ-region associated modules detected in rice with the same genetic backgrounds.

Table S4. The genes in the AZ-region associated modules detected in BHA and *aus* at each developmental stage.

Table S5. The genes in the AZ-region associated modules detected in SWR and *japonica* at each developmental stage.

Table S6. The hub genes in the AZ-region associated modules.

Table S7. The genes in the AZ-region associated modules in SWR only at each developmental stage.

Table S8. The enriched GO terms for the genes in the AZ-region associated modules detected in BHA and *aus* at each developmental stage.

Table S9. The enriched GO terms for the genes in the AZ-region associated modules detected in SWR and *japonica* or SWR only at each developmental stage.

Table S10. Genes related with lignin, cell wall and GA within each AZ-region associated modules.

Table S11. The genes overlapped in the AZ-region associated modules detected in SWR and *japonica* and BHA and *aus* at each developmental stage.

Table S12. The genes overlapped among AZ-region associated modules detected across different developmental stages in the same genomic backgrounds.

Table S13. Modules containing known seed shattering genes.

Table S14. The DE genes detected in BHA and *aus* at each developmental stage.

Table S15. The DE genes detected in SWR and *japonica* at each developmental stage.

Table S16. The AZ-exclusively up -or down regulated DE genes between weedy and cultivated rice.

Table S17. The overlapped AZ-exclusive DE genes between two weedy-cultivated rice comparisons.

Table S18. The enriched GO terms for the genes in the at each developmental stage.

Table S19. AZ-exclusive DE genes related with lignin, cell wall, GA and cell death.

Figure S1. Shattering trajectory for each weedy and cultivated genotype throughout development.

Figure S2. Clustering of RNA-seq samples from the *aus* lineage.

Figure S3. Clustering of RNA-seq samples from the *japonica* lineage.

Figure S4. Functional categories enriched in the genes within the AZ-region associated modules identified in the *aus* lineage at each developmental stage.

Figure S5. Functional categories enriched in the genes within the AZ-region associated modules identified in the *japonica* lineage at each developmental stage.

Figure S6. WGCNA modules containing known seed shattering genes.

