## Supplementary_figures for "Comparative tissue-specific transcriptomics reveal the genetic bases underlying evolutionary convergence of seed shattering in two independently evolved weedy rice lineages"

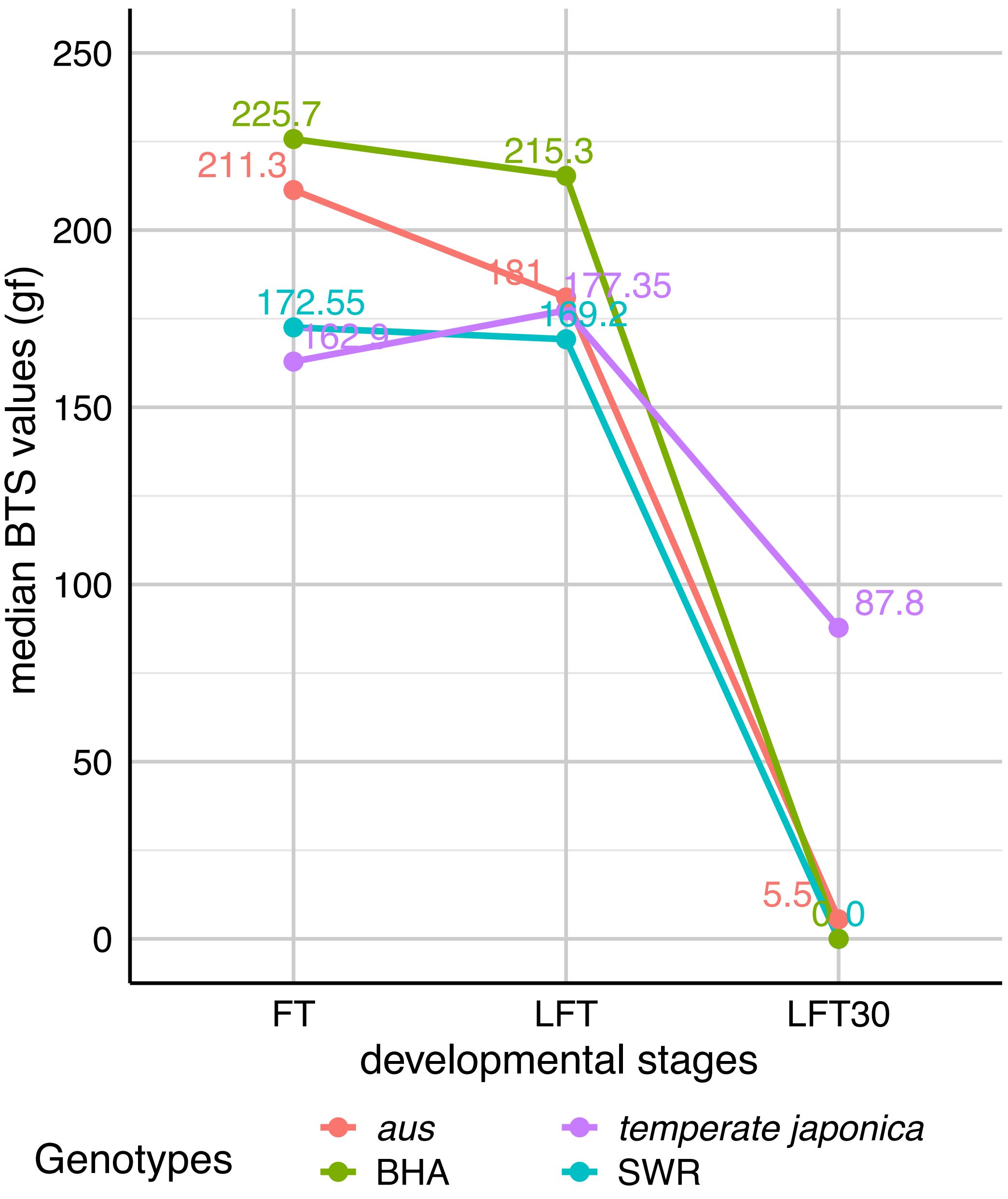


**Figure S1. Shattering trajectory for each weedy and cultivated genotype throughout development.**

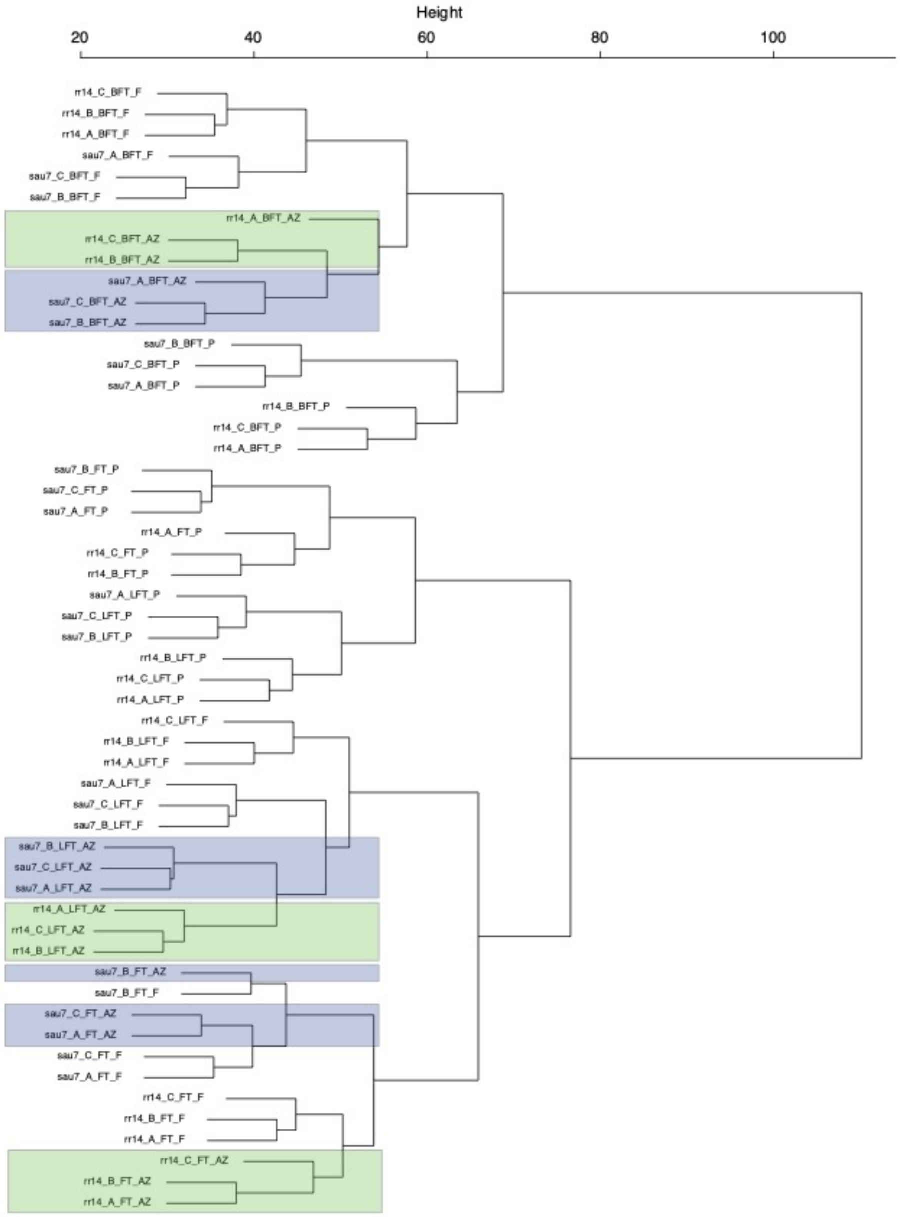


**Figure S2. Clustering of RNA-seq samples from the *aus* lineage.** The samples from the AZ regions of *aus* were highlighted as blue while the samples from the AZ regions of BHA were highlighted as green.

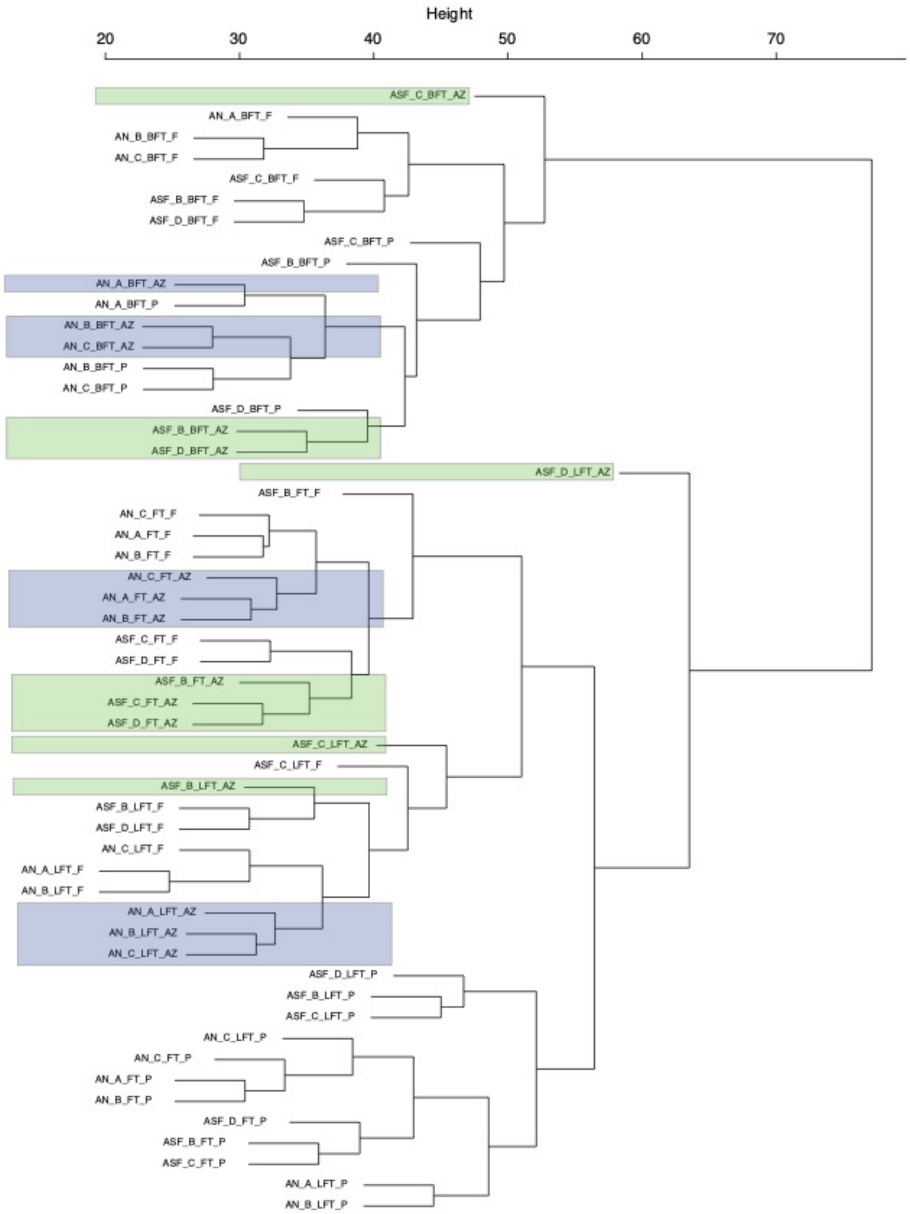


**Figure S3. Clustering of RNA-seq samples from the *japonica* lineage.** The samples from the AZ regions of *temperate japonica* were highlighted as blue while the samples from the AZ regions of SWR were highlighted as green.

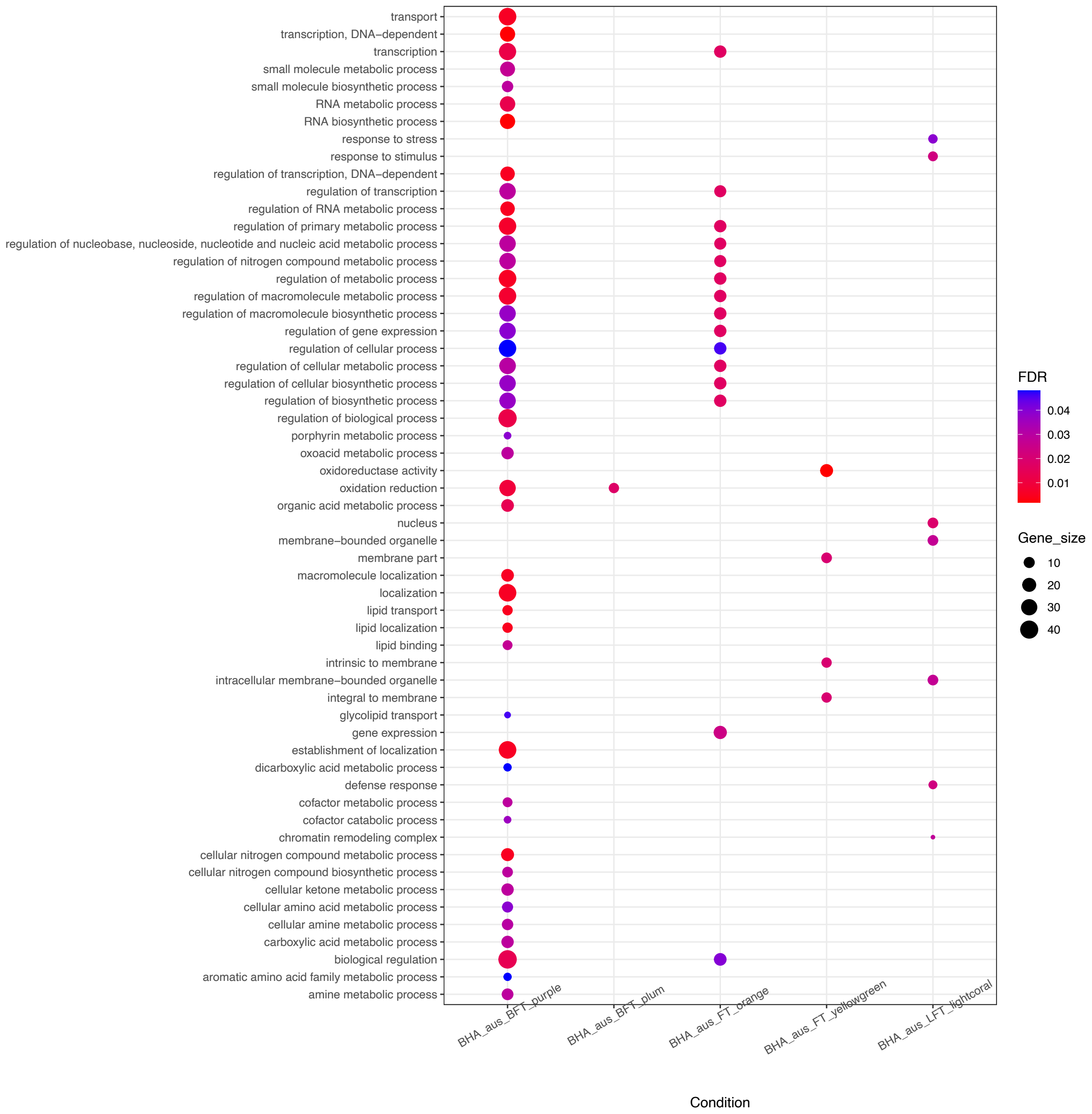


**Figure S4. Functional categories enriched in the genes within the AZ-region associated modules identified in the *aus* lineage at each developmental stage.** Significant Gene Ontology (GO) terms enriched in the genes within the AZ-region associated modules identified in the *aus* lineage at each developmental stage. Red and blue represent smaller and larger FDR values, respectively. The size of the dots represents the number of genes in each GO term over the total number of genes within the AZ-region associated module.

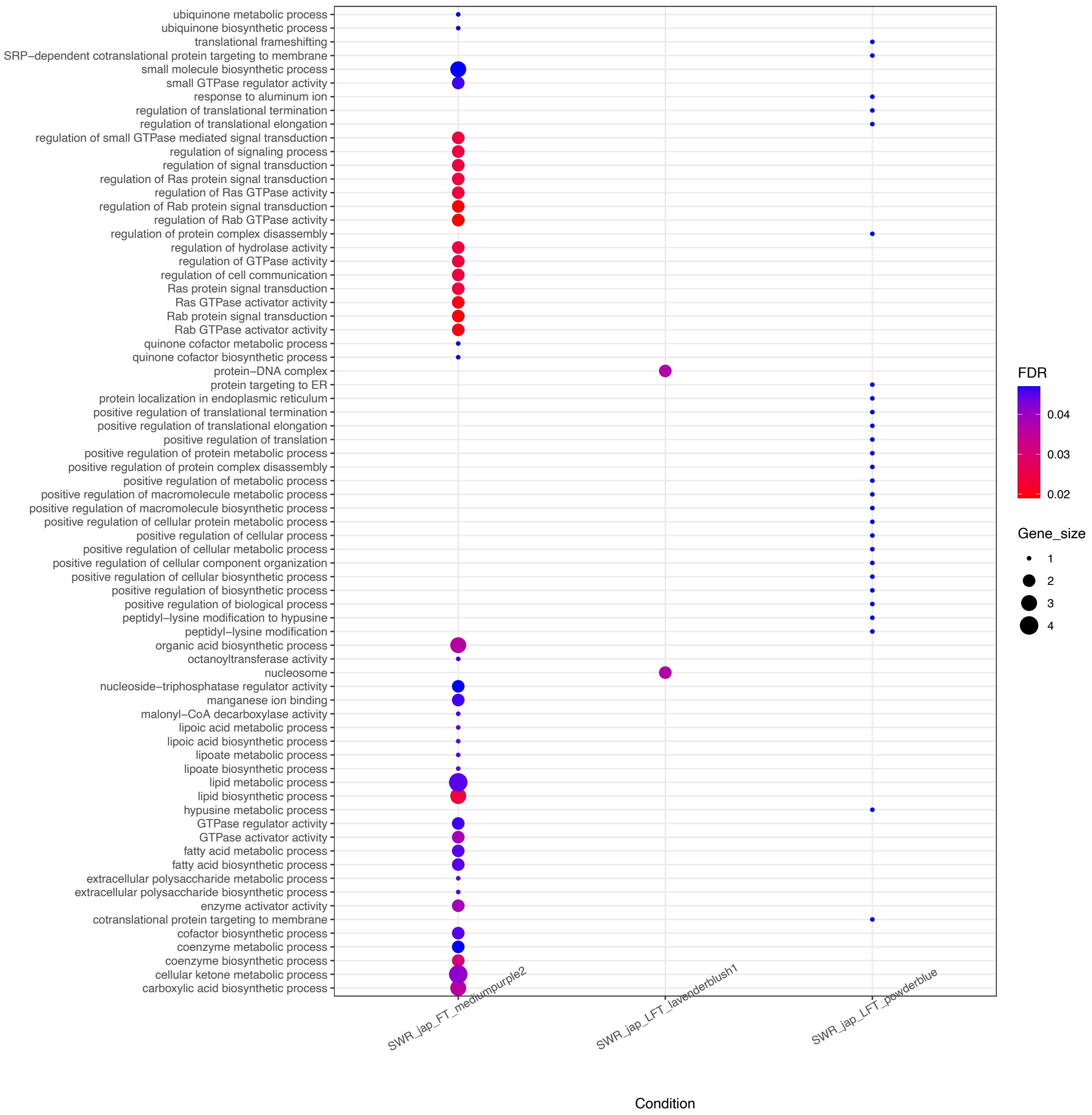


**Figure S5. Functional categories enriched in the genes within the AZ-region associated modules identified in the *japonica* lineage at each developmental stage.** Significant Gene Ontology (GO) terms enriched in the genes within the AZ-region associated modules identified in the *japonica* lineage at each developmental stage. Red and blue represent smaller and larger FDR values, respectively. The size of the dots represents the number of genes in each GO term over the total number of genes within the AZ-region associated module.

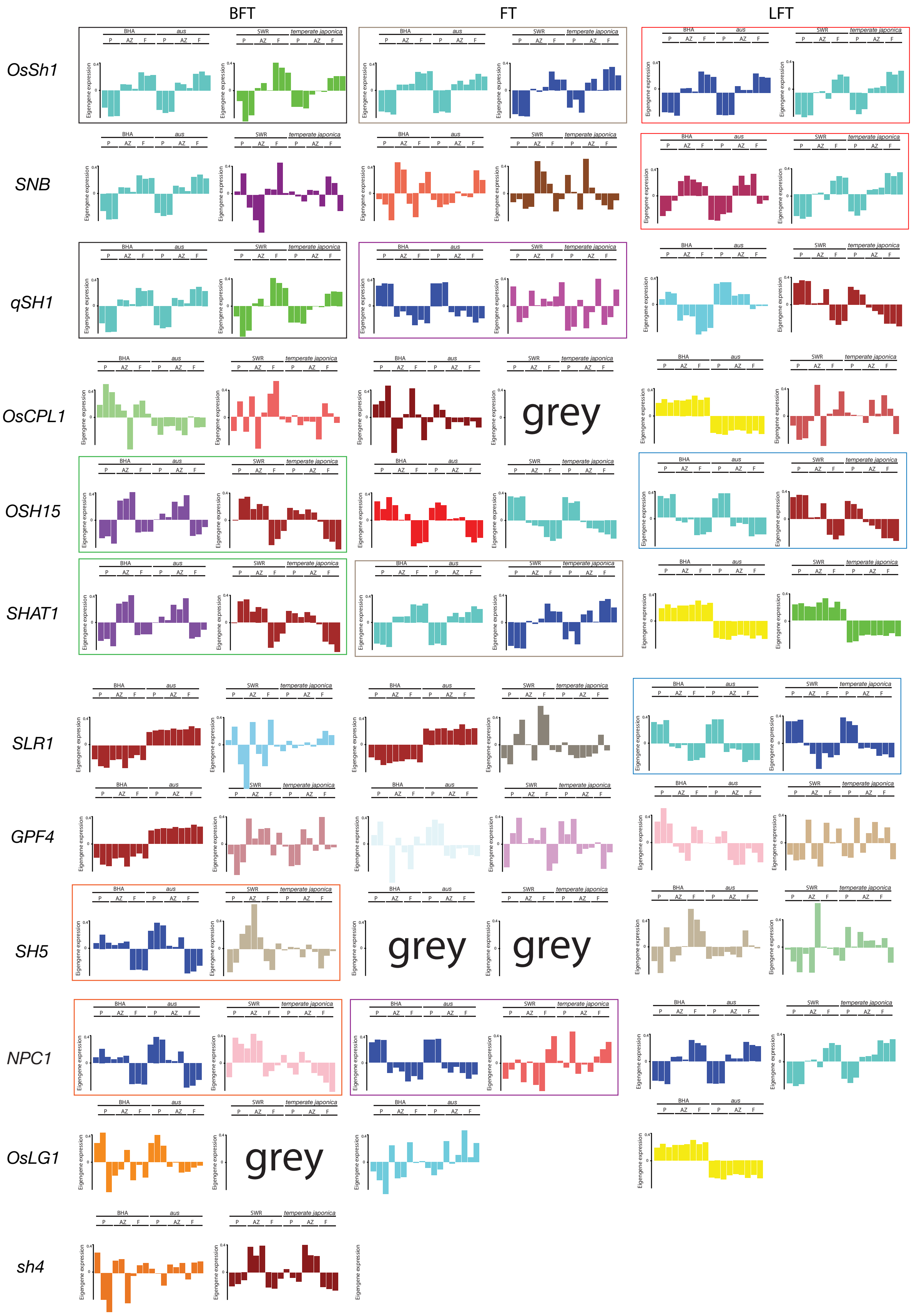
BFT
FT
LFT
OsSh1
SNB
qSH1
OsCPL1
OSH15
SHAT1
SLR1
GPF4
SH5
NPC1
OsLG1
sh4
BHA
aus
SWR
temperate japonica
P
AZ
F
Eigengene expression
0.4
0
grey

**Figure 6. WGCNA modules containing known seed shattering genes.** Eigengene expression boxplot for modules including the known seed shattering genes across different developmental stage. Grey module is the module containing all the unassigned genes.
